# Protracted speciation under the state-dependent speciation and extinction approach

**DOI:** 10.1101/2021.06.29.450466

**Authors:** Xia Hua, Tyara Herdha, Conrad Burden

**Affiliations:** Mathematical Sciences Institute, Australian National University, Canberra ACT 0200 Australia

**Keywords:** Protracted birth-death process, SSE approach, speciation completion rate, micro-macro evolution

## Abstract

How long does speciation take? What are the speciation processes that generated a species group? The answers to these important questions in evolutionary biology lie in the genetic difference not only among species, but also among lineages within each species. With the advance of genome sequencing in non-model organisms and the statistical tools to improve accuracy in inferring evolutionary histories among recently diverged lineages, we now have the lineage-level trees to answer these questions. However, we do not yet have an analytical tool for inferring speciation processes from these trees. What is needed is a model of speciation processes that generates both the trees and species identities of extant lineages. The model should allow calculation of the probability that certain lineages belong to certain species and have an evolutionary history consistent with the tree. Here we propose such a model and test the model performance on both simulated data and real data. We show that maximum likelihood estimates of the model are highly accurate and give estimates from real data that generate patterns consistent with observations. We discuss how to extend the model to account for different rates and types of speciation processes across lineages in a species group. By linking evolutionary processes on lineage level to species level, the model provides a new phylogenetic approach to study not just when speciation happened, but how speciation happened.

---

Speciation takes time (Avise 1999), yet most macroevolutionary models for speciation and extinction assume that speciation takes place instantaneously (Ricklefs 2007; Maddison et al. 2007). One exception is the protracted birth death model proposed by Etienne and Rosindell (2012), PBD hereafter, where a newly arising lineage takes time to become reproductively isolated from its ancestral species. The new lineage can be thought of a population newly isolated from the other populations of the same species. It is regarded as in an ‘incipient’ state before the completion of reproductive isolation and in a ‘good’ state after the completion. Such a model that accounts for different stages of speciation is critical to link microevolution on a lineage level to macroevolution on a species level, for two main reasons. First, it provides an important prior on the time window when gene flow or hybridization is possible between two lineages in a phylogenetic network (Huson et al. 2010). Second, it provides an alternative methodology for studying speciation, moving from studying sister species that are usually model organisms, to studying shared patterns in speciation processes across lineages in a species group (Network 2012).

PBD (Etienne and Rosindell 2012) considers a species-level tree that includes one representative lineage for each species. Fitting PBD to the tree allows us to estimate how often a new lineage arises (speciation initiation rate), how fast it develops reproductive isolation (speciation completion rate), and how often lineages go extinct (extinction rate). Lambert et al. (2015) derived an approximated likelihood function for PBD. The likelihood function is an approximation because it assumes that the representative lineage of a species in the tree is either the good lineage of the species, or the first descendant of the good but now extinct lineage of the species (Etienne et al. 2014). This approximation does not cause large bias in the estimates (Simonet et al. 2018).

However, fitting the model to phylogenies that only include one lineage of a species does not give accurate estimates of speciation completion rate from small phylogenies (< 400 species; Fig. S4 in Simonet et al. 2018), and the estimates of speciation initiation rate and extinction rate are unidentifiable (Simonet et al. 2018). This problem is common to many macroevolutionary models for speciation and extinction (Louca and Pennell 2020). One way to improve the accuracy of the protracted speciation model is to include incipient lineages in the tree, so that there are more lineages near the present, which carry most information on the extinction rate independent of speciation rate (Louca and Pennell 2020). Evolutionary histories among incipient lineages of the same species also carry important information on the speciation completion rate. The likelihood function by Lambert et al. (2015) can be applied to lineage-level trees, but it assumes trees as coalescent point processes, where node depths are a sequence of independent, identically distributed random variables. So, the likelihood function cannot account for the states (incipient or good) of extant lineages, because speciation processes depend on the states (e.g., speciation completion events cannot occur along an extant incipient lineage, because it will turn the lineage to a good state at present), which makes the tree no longer a coalescent point process (Lambert and Stadler 2013). For the same reason, the likelihood function cannot be extended to allow variation in the types or the rates of speciation processes across lineages. Therefore, we need a new model to describe the protracted speciation process, which not only accounts for the states (incipient or good) of extant lineages to improve estimation of speciation initiation rate, extinction rate, and speciation completion rate, but is also flexible enough to allow variation in speciation processes across lineages.

In this study, we provide the exact likelihood function of a new protracted speciation model applicable to both lineage-level and species-level trees. The calculation of the likelihood uses the state-dependent speciation and extinction approach (SSE; Maddison et al. 2007), so it is readily extended to account for variation in speciation processes across lineages (see Discussion). We call the new model protracted SSE or ProSSE. We assess the performance of ProSSE on simulated trees and compare the performance of ProSSE and PBD. We further demonstrate ProSSE using Australian rainbow skinks as a case study.

## MATERIALS AND METHODS

The difficulty in applying the SSE approach to PBD is the ambiguity in the definition of a good state, as the SSE approach requires the state of each extant lineage in the tree. In PBD, a good lineage is different from an incipient lineage in that a good lineage cannot become a distinct species from its ancestral species. But in reality, all lineages (i.e., populations) of a species have the potential to become a distinct species from the other lineages of the species, so all lineages are in the incipient state by definition. Even for a species that has only one lineage over time, anagenesis could still happen. For example, under the Bateson-Dobzhansky-Muller incompatibility model, the speciation completion rate is determined by how fast a population accumulates nearly neutral substitutions on loci that may cause incompatibility (Gavrilets 2014). When there is only one population of a species, the population can still accumulate substitutions on these loci and become a distinct species to its ancestral species, except that its speciation probability is one half of that between two populations, as either of the two populations can become a distinct species relative to the ancestral species.

In ProSSE, we redefine the good state as a ‘representative’ state, meaning that the lineage represents the species in the tree, so that each species has one representative lineage and multiple incipient lineages if any (Fig. 1a). Different from the good state, a representative lineage has the same evolutionary dynamics as an incipient lineage, i.e., they have the same speciation initiation rate (*b*), speciation completion rate (*λ*), and extinction rate (*μ*). The representative state is included only to indicate how many distinct species there are at a time slice in the tree. As we show below, no matter which lineage we pick as the representative of a species, the likelihood does not change. For example, in Fig. 1a, any of the three lineages of species A can be its representative lineage. In other words, we don’t need to assign a state to each lineage of the same species (and so dashed edges in Fig. 1b). This allows us to use the SSE approach to calculate the likelihood of the protracted speciation model, or the probability of a tree under the protracted speciation model.

**Figure 1.**
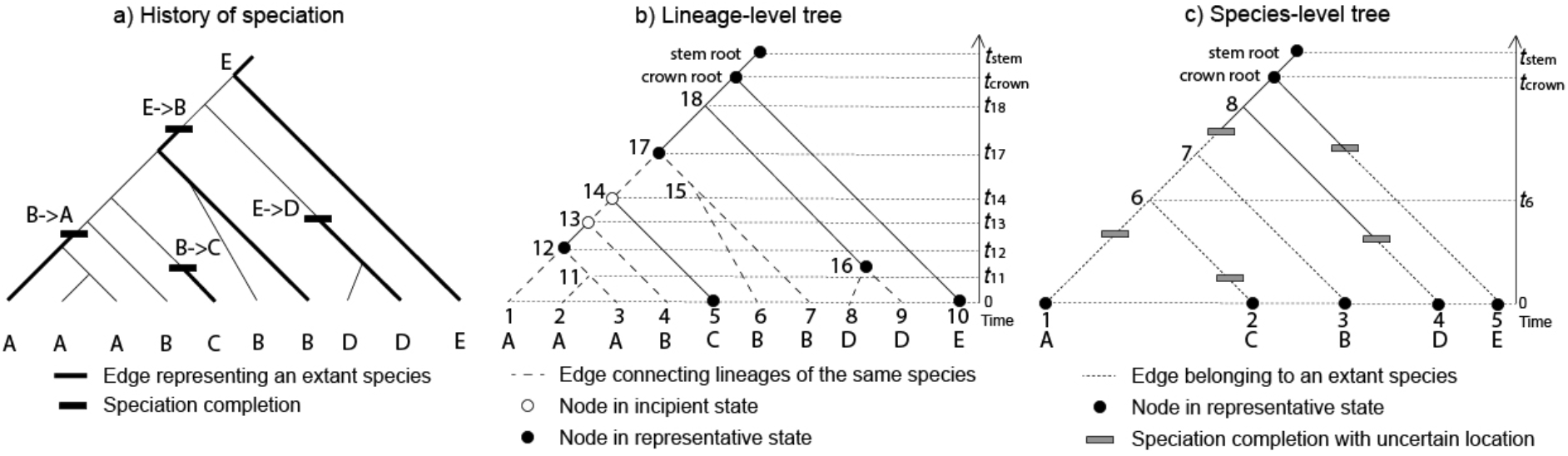
Illustration of ProSSE algorithms. From either lineage-level tree (b) or species-level tree (c), we don’t know exactly where speciation completion events occurred on the tree, but we can still get some information on the history of speciation (a) from these trees. From lineage-level tree (b), we know which lineages belong to the same species (dashed edges) and what state certain nodes have (circles). We are also sure that the speciation completion event that leads to an extant species must occur along an edge with ancestral node in incipient state and descendent node in representative state. For species-level tree (c), we only know that edges after the speciation completion event of an extant species should not leave any species that are different from all the extant species in the tree, but the location of the speciation completion event on the tree is uncertain.

### ProSSE algorithm for lineage-level trees

In common with other SSE models, ProSSE starts with states of the extant lineages in the tree, then, for each tip edge of the tree, calculates the probability that the edge stays as a single edge along the tree by integrating over all possible events (speciation initiation, speciation completion, and extinction) at any time point along the edge, calculates the joint probability of observing two sister tip edges at the node that connects them, and repeats the process along the internal edges and nodes until reaching the root to get the probability of the tree. What is different from other SSE models is that ProSSE uses different differential equations along edges connecting lineages of the same species (dashed edges in Fig. 1b) and edges connecting lineages of different species (solid edges in Fig. 1b). Along edges connecting lineages of the same species, for example, the dashed edges connecting the three A extant lineages in Fig. 1b, any speciation completion event would result in zero probability of observing these three A lineages in the tree. In contrast, suppose we have *N* distinct species at present, then at least *N* − 1 speciation completion events must have occurred along edges connecting lineages of different species (Fig. 1a).

Let’s define *D*_*R*_(*t*) and *D*_*I*_(*t*) as the probabilities that an edge in representative state, state *R*, and in incipient state, state *I*, at time *t* leads to extant descendants as observed in the tree. Here, *t* is measured backwards in time from the present at *t* = 0. Alternatively, we can think of *D*_*R*_(*t*) (resp. *D*_*I*_(*t*)) as the probability that an edge in state *R* (resp. state *I*) at time *t* stays as a single edge until its descendent node on the tree, and the boundary condition at the descendent node is determined by the probability of observing the part of the tree descending from that node. Also define *E*(*t*) as the probability that an edge at time *t* does not leave any descendant at present. Since *R* and *I* lineages have the same evolutionary dynamics, *E*(*t*) is the same for both. So, for all the edges in the tree, *E*(*t*) is calculated under the standard birth death process (Stadler 2010), with boundary condition *E*(0) = 0 and solution 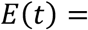 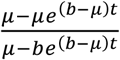

Along edges that connect lineages of the same species (dashed edges in Fig. 1b), *D*_*R*_(*t*) and *D*_*I*_(*t*) can be derived from

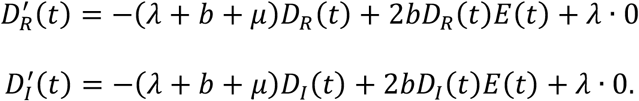

These equations are similar to the equations in BiSSE (Maddison et al. 2007), except that the last term on the r.h.s equals zero. This is because if a speciation completion event happens, then not all the lineages belong to the same species. As a result, a speciation completion event only reduces the probability of observing these edges connecting lineages of the same species, in both *D*_*R*_(*t*) and *D*_*I*_(*t*), as reflected in the first term. Note that these equations satisfy the same differential equation, but the initial condition is either *D*_*I*_(0) = 0, *D*_*R*_(0) = 1 for an *R* lineage, in which case *D*_*I*_(*t*) ≡ 0; or *D*_*I*_(0) = 1, *D*_*R*_(0) = 0 for an *I* lineage, in which case *D*_*R*_(*t*) ≡ 0. So to simplify the notation, define *D*(*t*) = *D*_*R*_(*t*) + *D*_*I*_(*t*), then

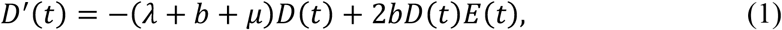

with the initial condition *D*(0) = 1. As a result, it is not necessary to assign a state to any edge that connects lineages of the same species. The solution at the ancestral node of an edge at time *t* + *s* is:

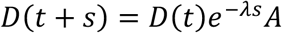

where *D*(*t*) is at the descendant node of the edge at time *t*, *s* is the finite edge length, and 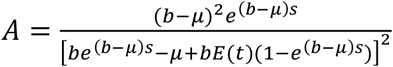.

Along edges connecting lineages of different species (solid edges in Fig. 1b), *R* and *I* lineages have different dynamics, because a speciation completion event resulting in an *R* state may occur in both cases. *D*_*R*_(*t*) and *D*_*I*_(*t*) can be derived from

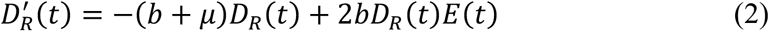

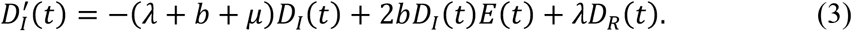

A speciation completion event has no effect on *D*_*R*_(*t*) in the equation because although an *R* lineage becomes a distinct species, it is still in *R* state, albeit representing a different species. The solutions at the ancestral node of an edge at time *t* + *s* are

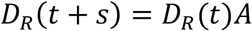

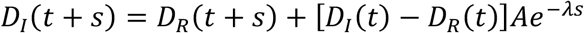

where *D*_*R*_(*t*) and *D*_*I*_(*t*) are at the descendant node of the edge at time *t* and *s* is the finite edge length. For an extant species with only one lineage, such as species C in Fig. 1b, its tip edge is also considered as connecting lineages of different species, so we apply equation 2 and 3 to the edge with initial conditions *D*_*R*_(0) = 1 and *D*_*I*_(0) = 0.

Consider the tree in Fig. 1b, where there are 10 extant lineages belonging to 5 distinct extant species. ProSSE first calculates from tip lineages that belong to the same species towards their common ancestors on the tree. For example, tip edges 1, 2, 3 belong to species A (Fig. 1b), so we calculate *D*(*t*) along these tip edges using Equation 1. Then, for each node along the edges connecting these tips, such as node 11, we calculate

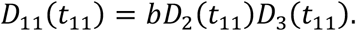

Here, we use the subscript to indicate which edge we are working on, for example, *D*_11_(*t*_11_) is the boundary condition at the descendant node of the edge linking node 11 and node 12 and the time at node 11 is *t*_11_. Repeating these calculations till node 12, we have *D*_12_(*t*_12_).

Next, ProSSE works on the edges connecting lineages of different species, which include the edges connecting the common ancestors of different extant species (e.g., the edge connecting node 17 and 18), the edges linking the common ancestor of some extant species to a lineage of a different extant species that is paraphyletic (e.g., the edge connecting node 12 and node 13; species B is paraphyletic), as well as the tip edge of a species that has only one extant lineage (e.g., tip edge 5). Along these edges, we calculate *D*_*R*_(*t*) and *D*_*I*_(*t*) using equations 2 and 3.

For edges that connect the common ancestors of different extant species, the boundary conditions at each common ancestor, for example, node 17, are *D*_*R*,17_(*t*_17_) = *D*_17_(*t*_17_), *D*_*I*,17_(*t*_17_) = 0, because the edge linking node 17 and 18 is the only lineage representing species B at *t*_17_, so node 17 must be in *R* state (Fig. 1b). Then, we can calculate along these edges towards the root of the tree. For example, at node 18, we have

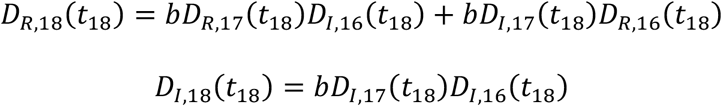

Note that these equations are different from BiSSE, because the new lineage generated from each speciation initiation event is always in the *I* state.

Edges connecting the common ancestor of some extant species to a lineage of a different extant species that is paraphyletic, for example, node 13 in Fig. 1b, are calculated as follows. One edge splitting from node 13 belongs to species B and the other edge leads to species A. Since the edge that leads to species A cannot represent species B, the state of the edge linking node 12 and node 13 at *t*_13_ must be in *I* state, so

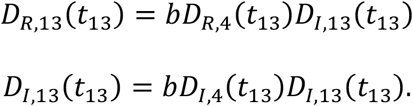

Since tip edge 4 belongs to species B, either *D*_*R*,4_(*t*_13_) = *D*_4_(*t*_13_) and *D*_*I*,4_(*t*_13_) = 0, or *D*_*R*,4_(*t*_13_) = 0 and *D*_*I*,4_(*t*_13_) = *D*_4_(*t*_13_), depending on the choice of *R* lineage for species B. Then, we calculate along the edge linking node 13 and 14, which is a dash edge (Fig. 1b), so the calculation uses equation 1 and the boundary condition at *t*_13_ is *D*_13_(*t*_13_) = *D*_*R*,13_(*t*_13_) + *D*_*I*,13_(*t*_13_).

Tip edges of a species that has only one extant lineage, for example, tip edge 5 in Fig. 1b, are calculated as follows. The calculation of node 14 is similar to node 13, so

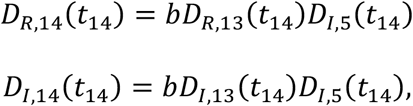

where *D*_*I*,5_(*t*_14_) is calculated using equations 2 and 3 with *D*_*R*,5_(0) = 1, *D*_*I*,5_(0) = 0. In this particular example, the edge linking node 14 and 17 is also a dashed edge belonging to species B, so *D*_14_(*t*_14_) = *D*_*R*,14_(*t*_14_) + *D*_*I*,14_(*t*_14_) = *b D*_13_(*t*_14_)*D*_*I*,5_(*t*_14_).

Last, we are at the crown root. Since there must be only one lineage representing the root species, so the crown root is in *R* state and the likelihood for the tree is *D*_*R,crown*_(*t*_*crown*_). If the tree has a root edge as in Fig. 1b, then the likelihood is further calculated along the root edge using Equation 2 with boundary condition *D*_*R,crown*_(*t*_*crown*_). To condition the final likelihood on the survival of the tree that has at least two distinct extant species, the likelihood is divided by *b*[1 − *E*(*t*_*stem*_)][1 − *E*_*R*_(*t*_*crown*_)], where *t*_*stem*_ = *t*_*crown*_ if the tree does not include a root edge and *E*_*R*_(*t*) is the probability that a lineage of a certain species at time *t* (root species at *t*_*crown*_ in this case) left no extant species that is different from its species identity. *E*_*R*_(*t*) is derived, as in Lambert et al. (2015), from

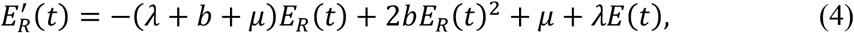

with boundary condition *E*_*R*_(0) = 1. The last term on the r.h.s. models a speciation completion event that leads to a distinct species but goes extinct before the present.

### ProSSE algorithm for species-level trees

The above ProSSE algorithm can be adapted to species-level trees that PBD uses, i.e., trees including one representative lineage of each extant species (see Fig. 1c). By representative sampling, we don’t know how many unsampled lineages there are in each extant species or how these lineages are placed in the tree. As a result, we can’t be sure of the species identity of any edge on the tree. Species identity is particularly important in this case. Let’s demonstrate this using the tree in Fig. 1c and assuming that we know where the speciation completion event that leads to each extant species occurred on the tree, so that we know which part of each edge belong to the same species as an extant species. Along a part of an edge that does not belong to any extant species (the solid part of the edges in Fig. 1c), we are certain that any lineage splitting from the part of the edge (except for the descendent node) went extinct before the present, so we can use *E*(*t*) to describe this probability as in equations 2 and 3. However, along a part of an edge that belongs to an extant species (the dotted part of the edges in Fig. 1c), we are only certain that any lineage splitting from the part of the edge (except for the descendent node) did not leave an extant species that is different from the species identity of the edge, so we need to use *E*_*R*_(*t*), instead of *E*(*t*), to describe this probability. For example, the edge linking node 6 and 7 in Fig. 1c can only leave extant lineages of species B.

Since we don’t know where speciation completion events occurred on the tree, we need to use different equations for edges belonging to an extant species and edges not belonging to any extant species, and use speciation completion rate as the transition rate between them. Whether an edge belonging to an extant species is denoted by the subscript after *R* or *I* state, with 1 for belonging to an extant species and 0 for not belonging to any extant species. For example, *D*_*R*1_(*t*) is the joint probability of the part of tree descended from an edge in state *R* at time *t* and the edge’s species identity being an extant species. These equations are

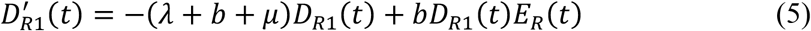

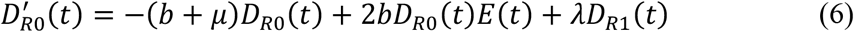

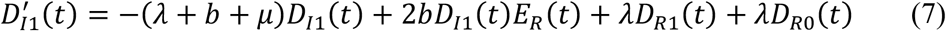

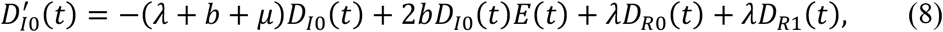

with the boundary condition for each tip edge at present as *D*_*R*1_(0) = 1, *D*_*R*0_(0) = 0, *D*_*I*1_(0) = 0, *D*_*I*0_(0) = 0. Note that *D*_*R*_(*t*) = *D*_*R*1_(*t*) + *D*_*R*0_(*t*) and *D*_*I*_(*t*) = *D*_*I*1_(*t*) + *D*_*I*0_(*t*), such that these equations equate to equations 2 and 3 when all lineages are sampled.

Equations 6 and 8 are for edges not belonging to any extant species. They are similar to equations 2 and 3, except that we need *λ D*_*R*1_(*t*) in both equations to describe the event where an edge in *R* or *I* state becomes an *R* lineage of an extant species. Equations 5 and 7 are for edges belonging to an extant species. Equation 5 is similar to equation 1, because a speciation completion event along an *R* lineage of an extant species will change its species identity and so will only reduce the probability of observing the lineage. Equation 5 differs from equation 1 in that 1) *E*(*t*) is replaced by *E*_*R*_(*t*) to account for any unsampled lineages of the same species; 2) the coefficient of the last term on the r.h.s changes from 2 to 1. The coefficient is 2 in equation 1 because the tree does not change when either edge splitting from a speciation initiation event is the observed edge, as the other edge must be extinct. This is no longer true in equation 5, because both edges may leave some extant lineages of the species and so the observed edge has to be the one that led to the sampled lineage of the species, otherwise the tree will be different. Similarly, we need to replace *E*(*t*) by *E*_*R*_(*t*) in equation 7 and add the term *λ D*_*R*0_(*t*) to describe the event where an *I* lineage of an extant species becomes a new species that does not leave any extant lineage of its species identity with probability *D*_*R*0_(*t*).

Equations 5–8 have no easily determined analytical solution, so we numerically integrate the equations along each edge. At each internal node, such as node 6 in Fig. 1c, we have

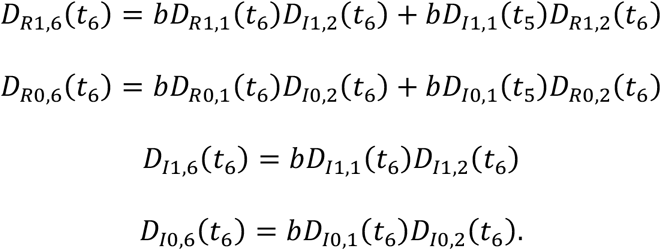

Repeating the calculation along each edge from tips to the crown root, or to the stem root if the tree has a root edge. The root must be in an *R* state, so the final likelihood is *D*_*R*1_(*t*_*stem*_) + *D*_*R*0_(*t*_*stem*_), where *t*_*stem*_ = *t*_*crown*_ if the tree has no root edge. To condition the likelihood on the survival of the tree, *D*_*R*1_(*t*_*stem*_) is divided by *b*[1 − *E*(*t*_*stem*_][1 − *E*_*R*_(*t*_*crown*_], but *D*_*R*0_(*t*_*stem*_) is divided by *b*[1 − *E*_*R*_(*t*_*crown*_][1 − *E*_*R*_(*t*_*crown*_)]. This is because if the root does not belong to any existing species, then both edges split from the crown root should leave some extant species that are distinct to the root species.

### Assessing ProSSE performance

We assess the performance of ProSSE by first simulating lineage-level trees under our protracted speciation model, then searching for the maximum likelihood (ML) estimates of each parameter in the model using the ProSSE algorithm for lineage-level trees, and last reporting the deviation in the ML estimates to the true parameter values used in the simulation. For each simulated tree, we also construct its species-level tree by randomly picking a representative lineage of each extant species in the tree and discarding the other lineages. We then search for the ML estimates of each parameter in the model using the ProSSE algorithm for species-level trees, report the deviation in the ML estimates to the true parameter values used in the simulation, and compare the deviation to that reported in Simonet et al. (2018) for PBD.

We simulate trees using the same parameter sets as used in Simonet et al. (2018). In brief, we simulate 1000 trees under our protracted speciation model for various combinations of parameter values (*b* = 0.3, 0.4, 0.5, 0.6, 0.7; *μ* = 0, 0.1, 0.2; *λ* = 0.1, 0.3, 1). But our simulation is different from the PBD simulation in Simonet et al. (2018) in three ways. First, we do not track the good or incipient state of each lineage over time but track each lineage’s species identity. Second, all the lineages in our simulation can go through speciation initiation, speciation completion, and extinction events. Third, Simonet et al. (2018) fixed the crown age of each simulated tree to 15, but this will exclude many possible trees we may observe in nature under high extinction rate. Instead, we fix the stem age of each simulated tree to 15. This difference will result in smaller and less informative trees in our simulation with high extinction rate, compared to the PBD simulation.

We implement the ProSSE algorithms and the simulation in the R package for SSE models: ‘diversitree’ (FitzJohn 2012). The implementation includes new R and C functions, new Rd files, and updated namespace file to the package. These are available at github.com/huaxia1985/ProSSE. Users can copy these files to the package source code, compile, and install the modified package in R. The main functions are make.prosse for the algorithm for lineage-level tree, make.prosse.sp for the algorithm for species-level tree, and make.tree.prosse for simulating trees under our protracted speciation model. These functions can be used in the same ways as the other SSE models in the package, for example estimating parameters using both ML and Bayesian approaches.

### Case study: Australian rainbow skinks

To test the performance of ProSSE on real species groups, we use Australian rainbow skinks as the case study. Rainbow skinks include three recognised genera: *Carlia*, *Lygisaurus* and *Liburnascincus*, which together contain 41 named species in Australia. Many of these species have clear phylogeographic lineages. Bragg et al. (2018) published a preliminary multispecies coalescent tree of the species group and Bragg et al. (unpublished) updated the tree with a complete sample of all recognized species and all phylogeographic lineages within each species using StarBEAST2 (Ogilvie et al. 2017). Singhal and Moritz (2013) found absence of introgression in the hybrid zone between *Carlia rubrigularis* N and S lineages in the group. This provides key evidence of reproductive isolation between the two lineages and *C. rubrigularis* N lineage is now elevated to a distinct species *C. crypta* (Singhal et al. 2018). Singhal et al. 2018 suggested that the divergence time between *C. crypta* and *C. rubrigularis* (*t*_*c*_) is a conservative estimate of the amount of time to complete speciation in the group, because two of the four pairs of sister species in two typical species groups (*C. rubrigularis* group and *Lampropholis coggeri* group) of the subfamily that contain the rainbow skinks have shorter divergence time than *t*:. The completeness of the lineage-level tree and independent evidence on the time to complete reproductive isolation make Australian rainbow skinks a good case study to test the performance of ProSSE. If ProSSE can reliably estimate speciation initiation rate *b*, extinction rate *μ*, and speciation completion rate *λ* in real species groups, then the estimated parameter values for Australian rainbow skinks should have high probability to result in two out of four pairs of sister species having shorter divergence time than *t*_*c*_, assuming that lineages in the species group have similar speciation processes (see Discussion).

Using ProSSE for lineage-level trees, we get the ML estimates of the three parameters from each of the 1800 posterior samples of the multispecies coalescent tree of the species group (Bragg et al. unpublished; Fig. 5), in order to account for phylogenetic uncertainty. For each sample of the tree with root depth *t*_*root*_ and the corresponding ML estimates of the three parameters, we derive the probability density distribution of divergence time between sister species in the species group and use this distribution to calculate the proportion of divergence time between sister species shorter than *t*_*c*_. To derive the probability density distribution, we apply ProSSE for species-level trees to calculate the probability of a tree (unconditional on the survival of the tree) that consists of one pair of sister species linked by a crown root with tip edge length *t*_*d*_. Since the tree is just a pair of sister species with divergence time *t*_*d*_, the tree probability unconditional on tree survival gives the probability of observing a pair of sister species with divergence time *t*_*d*_. Integrating the tree probability over *t*_*d*_ from 0 to *t*_*root*_ gives the overall probability of observing a pair of sister species. Then, the tree probability divided by the integral gives the probability density of *t*_*d*_.

## RESULTS

### High accuracy of ProSSE estimates for lineage-level trees

For speciation initiation rate *b* and extinction rate *μ*, the absolute medians of error in ProSSE ML estimates are close to zero (*b*: 0.01 ± 0.005; *μ*: 0.03 ± 0.028) over all simulation scenarios (Fig. 2a-e and Fig. 3a-e). The interquartile ranges of error decrease rapidly when the number of lineages increases (Fig. 2f and Fig. 3f), from *b*: −0.16~0.46 and *μ*: −0.16~0.84 for trees with 5 to 50 tip lineages, to *b*: −0.07~0.11 and *μ*: −0.10~0.19 for trees with over 50 tip lineages. These results suggest that ProSSE ML estimates of speciation initiation rate and extinction rate for completely sampled tree are median-unbiased consistent estimators, with high accuracy for trees on lineage level that typically consist of over 50 tip lineages.

**Figure 2.**
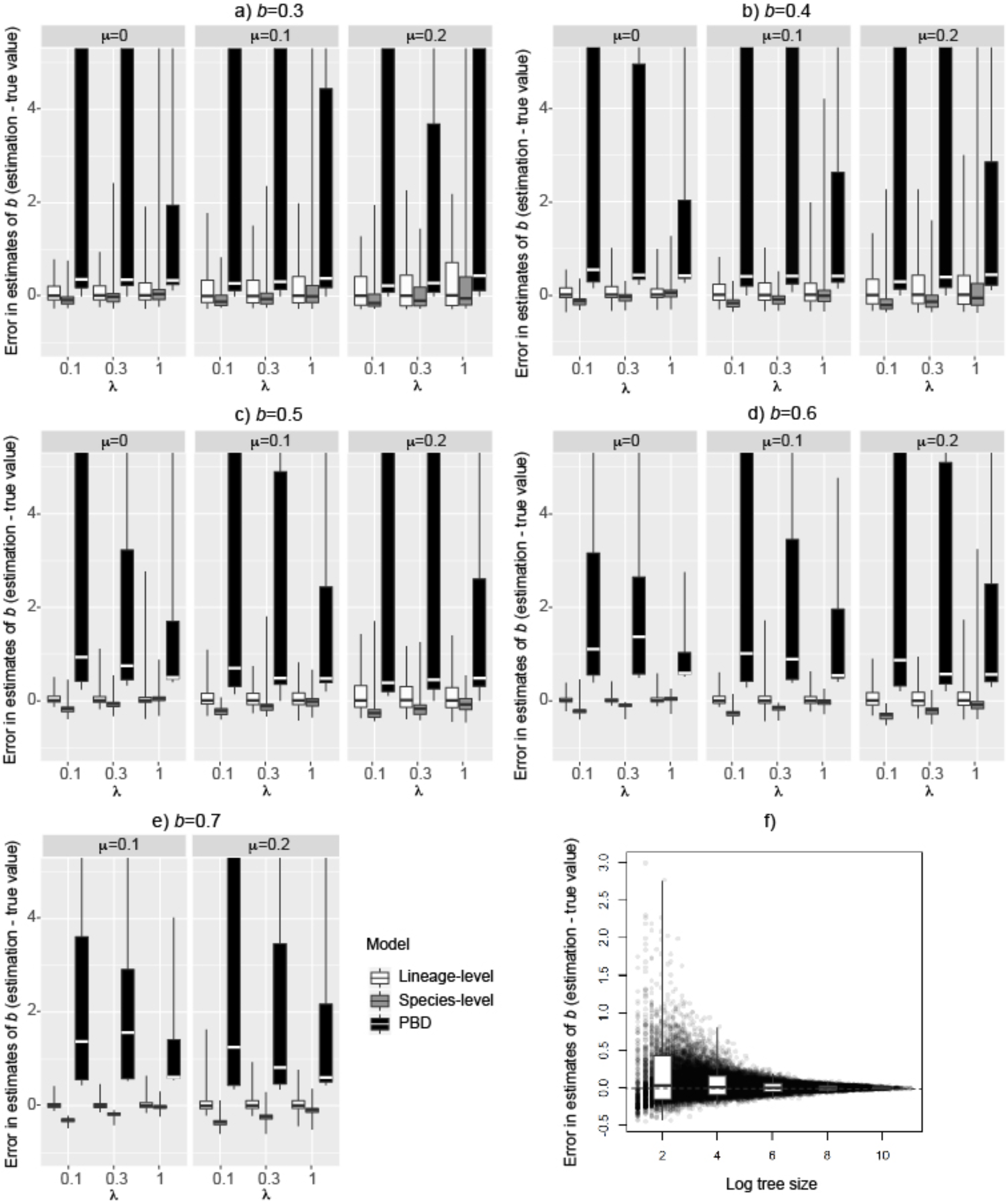
Error in ML estimates of the speciation initiation rate *b* for each simulation scenario. Each plot of a-e) represents a value of *b* used in the simulation. Each facet represents a value of *μ* used in the simulation. Each tick on x-axis represents a value of *λ* used in the simulation. Each boxplot represents the distribution of errors, the difference between the estimated and the true value of *b* over 1000 simulated trees, showing the minimum, the maximum, the median, the first and third quartiles of the distribution. In cases of extreme values, boxplots are cut off for graphical readability. White boxplots are errors of ProSSE estimate on lineage-level tree. Grey boxplots are errors of ProSSE estimate on species-level tree. Black boxplots are errors of PBD estimate on species-level tree with randomly sampled lineage as the representative lineage of each species. The data for PBD is from Simonet et al. (2018). f) plots errors of ProSSE estimate of *b* on lineage-level tree over all simulation scenarios against log tree size (the number of lineages in the simulated tree on log scale), with each datapoint corresponding to a simulated tree. White boxplots in f) show the distribution of errors for log tree size between 2 and 4, between 4 and 6, between 6 and 8, 8 and 10, and over 10.

**Figure 3.**
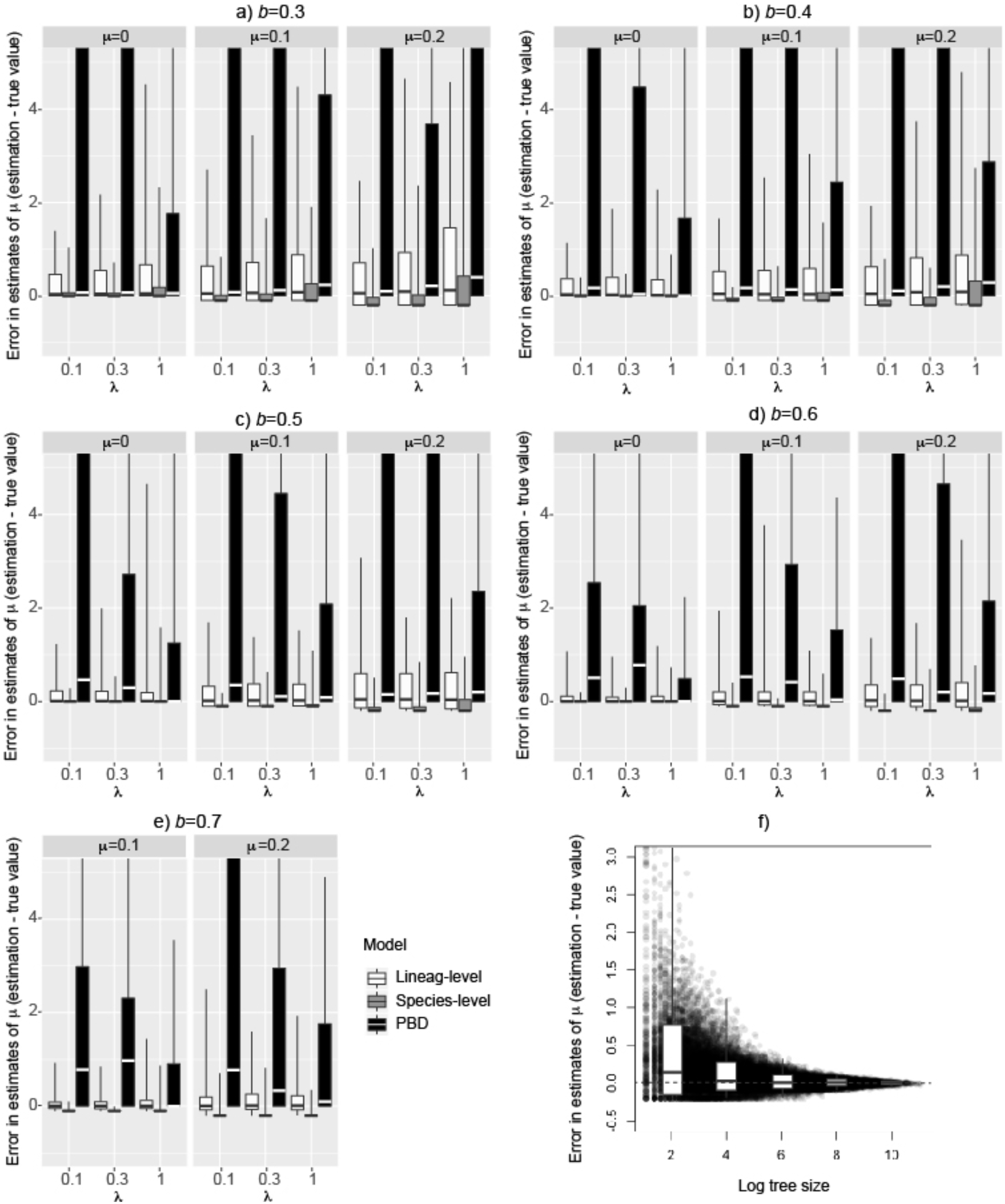
Error in ML estimates of the extinction rate *μ* for each simulation scenario (a-e) and the decrease of error bounds over tree size for ProSSE estimate of *μ* on lineage-level tree (f). See figure details in the legend of Figure 2.

For speciation completion rate *λ*, the absolute medians of error in ProSSE ML estimates are close to zero (0.01 ± 0.017) when *λ* ≤ *b* − *μ*, where *b* − *μ* is lineage diversification rate (Fig. 4a-e). When *λ* > *b* − *μ*, ProSSE overestimates *λ*, particularly for trees with all extant species having a single lineage, i.e., no extant lineage is in incipient state (compare boxplots in white and shaded with lines in Fig. 4a-e). These are the trees with little information on the upper bound of *λ*, because speciation completion only needs to happen fast enough to make all extant lineages become distinct species. In contrast, incipient lineages give information on the lower bound of *λ*, because speciation completion needs to happen sufficiently slowly, so that it does not happen along any incipient lineage. As a result, the proportion of lineages representing a single-lineage species in a tree determines whether ProSSE overestimates speciation completion rate.

**Figure 4.**
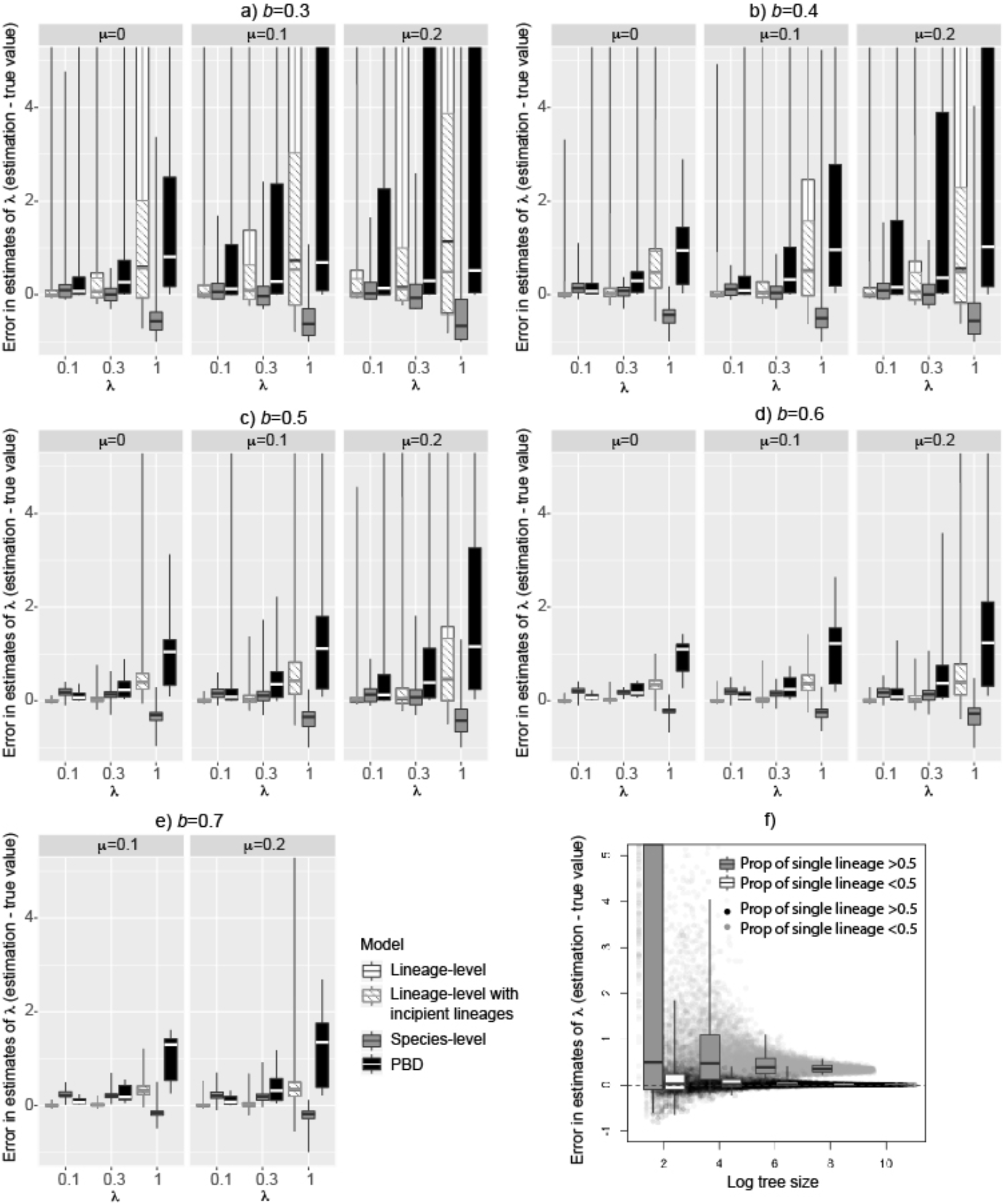
Error in ML estimates of the extinction rate *λ* for each simulation scenario (a-e) and the decrease of error bounds over tree size for ProSSE estimate of *λ* on lineage-level tree (f). ProSSE estimate of *λ* is biased for trees with more than half lineages representing single-lineage species, so (f) plots trees with under and above half lineages representing single-lineage species separately, with grey dots and grey boxplots for trees with more than half lineages being single-lineage species and black dots and white boxplots for trees with less than half lineages being single-lineage species. See figure details in the legend of Figure 2.

The threshold for this proportion above which ProSSE overestimates *λ* is about 0.5 (Fig. 3f): when tree size increases, errors in the ProSSE estimate for trees with over half lineages representing single-lineage species converges to 0.33, while errors for trees with under half lineages representing single-lineage species converge to zero. These results suggest that ProSSE ML estimate is an unbiased consistent estimator for speciation completion rate when trees have under half their lineages representing single-lineage species.

The interquartile range of error decreases rapidly from −0.11~0.26 for trees with 5 to 50 tip lineages, to −0.02~0.08 for trees with over 50 tip lineages. When trees have over half their lineages representing a single-lineage species, the ProSSE ML estimate is an inconsistent estimator with bias 0.33 and the interquartile range of error reduced with tree size, from −0.11~49 for trees with 5 to 50 tip lineages to 0.26~0.52 for trees over 50 tip lineages. Overall, ProSSE gives reasonably good estimate for speciation completion rate for lineage-level trees that typically consist of over 50 tip lineages.

### ProSSE outperforms PBD for species-level trees

For species-level trees, i.e., trees with only one representative lineage sampled for each species, both ProSSE and PBD give biased estimators for the three parameters (Fig. 1–3a-e). Over all simulation scenarios, ProSSE ML estimates have smaller biases (the absolute medians of error for *b*: 0.12 ± 0.090; *μ*: 0.11 ± 0.081; *λ*: 0.21 ± 0.168) than PBD ML estimates (*b*: 0.61 ± 0.330; *μ*: 0. 25 ± 0.239; *λ*:0.47 ± 0.432), as well as narrower interquartile range of errors (*b*: 0.17 ± 0.116; *μ*: 0.08 ± 0.138; *λ*:0.27 ± 0.171) than PBD (*b*: 16 ± 31.5; *μ*: 16 ± 31.5; *λ*:1.58 ± 2.190). In particular, speciation initiation rate *b* and extinction rate *μ* are generally unidentifiable in PBD. They are identifiable in ProSSE, but both are underestimated (Fig. 1–3a-e).

ProSSE tends to underestimate *b* when the speciation completion rate *λ* is lower. This is because more incipient lineages are unsampled in trees generated under low *λ*, which results in loss of information on recent speciation initiation events and so lower estimation of *b*. The underestimation of *μ* is consistent across simulation scenarios. This is a common problem due to incomplete sampling in the SSE approach (FitzJohn et al. 2009), because incomplete sampling reduces the effect of *μ* on the extinction probability *E*(*t*), e.g., in ProSSE, *E*(*t*) becomes *E*_*R*_(*t*), so the extinction rate affects tree likelihood mainly via the −*μ* term in equations 5 and 7, which makes the ML estimate of *μ* lower than the true extinction rate.

As a result of underestimation in the speciation initiation rate *b* under a lower speciation completion rate *λ* and consistent underestimation of the extinction rate *μ*, ProSSE underestimates the lineage diversification rate (*b* − *μ*) under low *λ* (*λ* = 0.1 in Fig. 4) and overestimates it under high *λ* (*λ* = 1 in Fig. 4). This leads to overestimation of *λ* when *λ* is small (under *λ* = 0.1 in Fig. 3a-e) and underestimation when *λ* is high (under *λ* = 1 in Fig. 3a-e), because an underestimated lineage diversification rate increases the probability that a tree has no unsampled incipient lineages, and so favours a high speciation completion rate, and vice versa. Compared to ProSSE, PBD consistently overestimates lineage diversification rate and speciation completion rate with larger biases.

### ProSSE gives reliable estimates from real data

For Australian rainbow skinks, ProSSE estimates, on average, 0.27 speciation initiation rate (Fig. 5b), nearly zero extinction rate (Fig. 5c), and 0.31 speciation completion rate (Fig. 5d). These estimates mean that, along a given lineage of Australian rainbow skinks, the interval between two successive speciation initiation events along the lineage follows an exponential distribution with mean ~3.7 Myr. After a speciation initiation event splits the lineage into two, the time for either of the two lineages to become a new species follows an exponential distribution with mean ~3.2 Myr. These estimates lead to nearly half extant sister species in the species group having shorter divergence time than that between *C. Crypta* and *C. rubrigularis*, which gives about 0.35 probability that two out of four pairs of extant sister species have shorter divergence time than that between *C. Crypta* and *C. rubrigularis* (Fig. 5e). This result agrees with Singhal et al. (2018) that the divergence time between *C. Crypta* and *C. rubrigularis* is a conservative cut-off to delimit distinct species in Australian rainbow skinks, suggesting that ProSSE is able to give reasonable estimates of parameters from real data.

**Figure 5.**
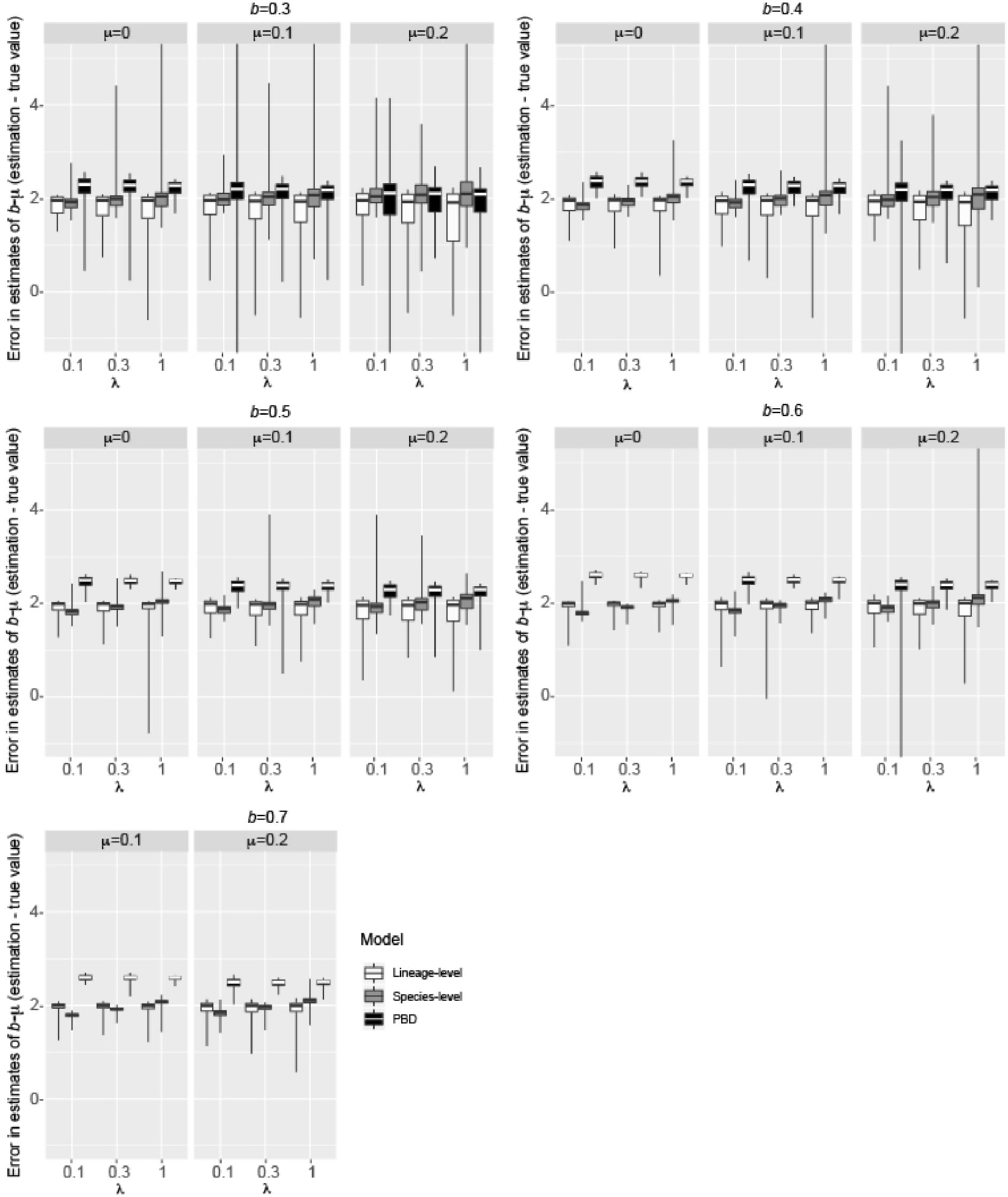
Error in ML estimates of lineage diversification rate (*b* − *μ*) for each simulation scenario. See figure details in the legend of Figure 2.

**Figure 6.**
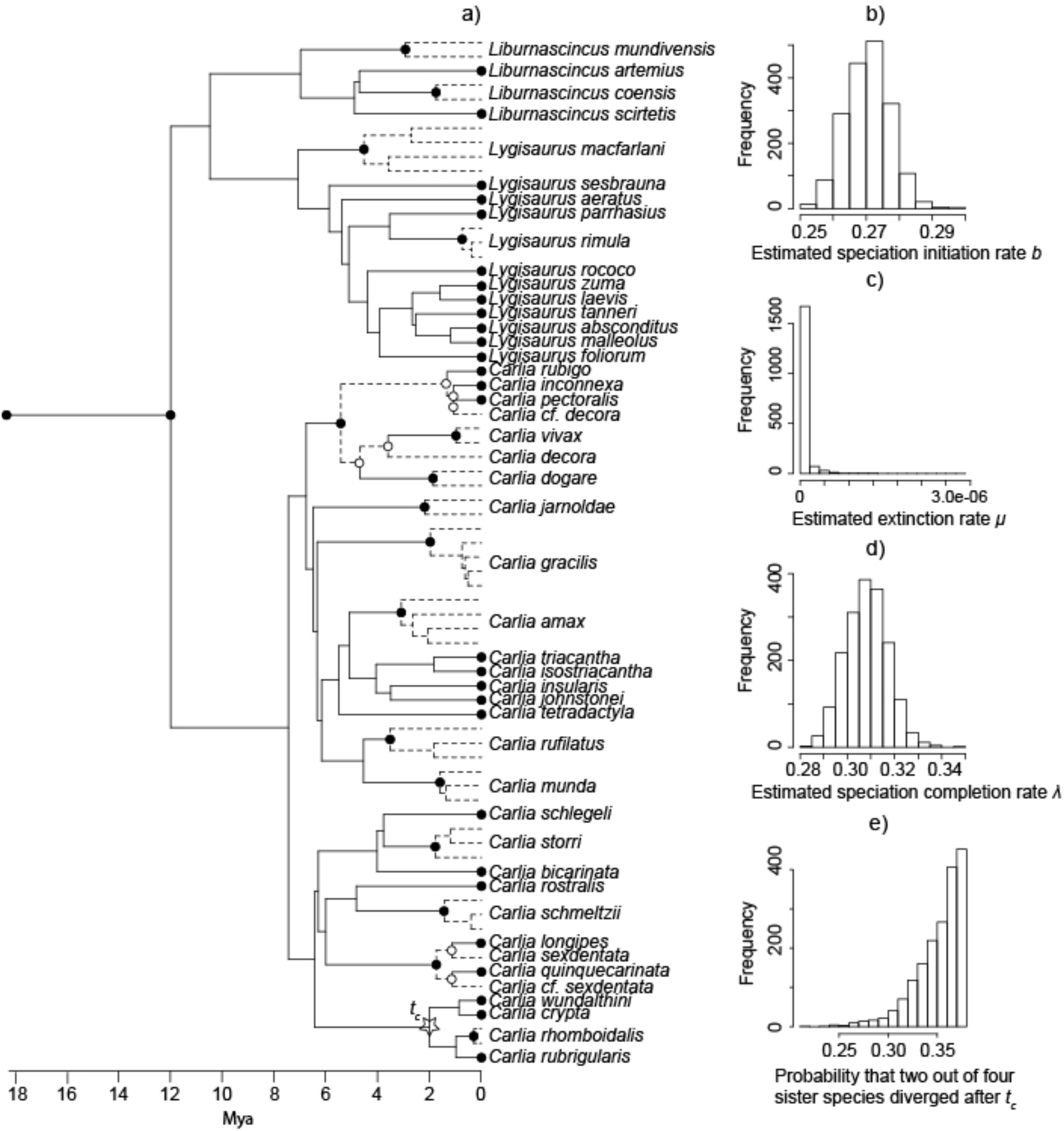
Maximum clade credibility tree of Australian rainbow skinks on lineage level (a) and the results of ProSSE analysis on the tree (b-d). The information we know from the tree is marked in (a), with dashed edges belong to the same species, nodes with black circles in representative state, and nodes with white circles in incipient state. Plots (b-d) show the distribution of ProSSE ML estimates for speciation initiation rate (b), extinction rate (c), and speciation completion rate (d) over 1800 posterior samples of the tree. Under these sets of parameter values, (e) plots the probability that two out of four randomly sampled pairs of sister species have divergence time lower than that between *C. crypta* and *C. rubrigularis* (*t*_*c*_, the node height indicated by a star in (a).

## DISCUSSION

This study provides an exact likelihood function for the protracted speciation model, using state dependent speciation extinction approach (ProSSE). We show that, for completely sampled trees on lineage level, ProSSE is able to give accurate estimates for trees of typical size in macroevolutionary analyses. When applied to real data, ProSSE is able to give reliable estimates that are consistent with independent evidence on species divergence time. For species-level trees with representative sampling, ProSSE ML estimates have consistently lower biases and narrower bounds of errors than PBD ML estimates for the three parameters. In summary, ProSSE gives good estimates for the three parameters over a wide range of conditions.

ProSSE has three main assumptions. The first assumption, complete sampling for lineage-level trees, is relatively minor as the assumption can easily be relaxed in a similar way to how we account for representative sampling in species-level trees. The probability that a lineage of species *i* is not included in the tree can be described by *E*_*i*_(*t*) using the same equation as for *E*(*t*), except that the initial condition is not zero but the fraction of unsampled lineage of species *i* (FitzJohn et al. 2009). For species with different sampling fractions, we need to use separate equations for lineages belonging to these different species, same as we have done in equations 5–8 to treat lineages not belonging to any extant species and lineages belonging to an extant species separately. So at the root of the tree, we will have the joint probabilities of the tree and the root belonging to a certain extant species, in addition to the joint probability of the tree and the root not belonging to any extant species, as we did for species-level trees.

The two remaining assumptions of ProSSE are 1) that the species identity of each extant lineage is known; and 2) that all lineages in the species group have similar rates and types of speciation processes. These two assumptions do not hold for many species groups. But since ProSSE treats each edge in a tree separately, we are able to relax these assumptions under the same model framework. Below, we discuss how to extend ProSSE to relax these assumptions.

### Simultaneous inference for species identity and speciation process

ProSSE assumes that the species identity of each extant lineage is known. We can relax this assumption by simultaneously inferring species identity and parameters in ProSSE. Given two lineages which may or may not belong to distinct species, we can introduce a binary variable at the most recent node connecting the two lineages in the tree with value 1 if the two lineages belong to distinct species and 0 otherwise. Applying this to any two lineages with uncertain species identity, we construct a set of binary variables, where each variable is associated with a node. Then a Markov chain Monte Carlo (MCMC) algorithm can be applied to approximate the joint probability distribution of the ProSSE parameters and these binary variables. This method is similar to MCMC methods for variable selection (Chipman et al. 2001). Since the probability of each binary variable conditional on the other binary variables and the ProSSE parameters can be calculated analytically from Bayes’ theorem and the ProSSE likelihood function, Gibbs sampling can be applied to each binary variable sequentially, while ProSSE parameters are sampled by a Metropolis-Hastings algorithm. As a result, the number of posterior samples with the binary variable of a node equal to one gives the probability that the two extant lineages connected by the node belong to distinct species. This will allow use of lineages of known species identity to inform ProSSE parameters and to delimit species in the tree for lineages with uncertain species identity.

### Accounting for different speciation processes in the tree

Another critical assumption of ProSSE is that all lineages in the species group have similar rates and types of speciation processes, so that the same set of ProSSE parameter values is fitted to all lineages. ProSSE can be extended to relax this assumption, as the SSE approach is designed to account for variation in speciation and extinction rates among lineages (Maddison et al. 2007). For example, although the majority of Australian rainbow skinks are generalists and distributed in the Australian tropical savanna, there are still some species distributed in rainforests, potentially having undergone allopatric speciation in glacial refugia (Graham et al. 2006), and some other adapted to rock habitat that potentially have undergone ecological speciation (Blom et al. 2016). Since the relative prevalence of different speciation processes is largely associated with habitats, we can use the habitat of extant lineages to inform variation in speciation processes among lineages.

For example, there are a large number of cryptic lizard species in the Australian rainforests (Moritz et al. 2009), suggesting that most speciation in the area is completed by the accumulation of incompatible genes in allopatry. If this is true, then the speciation completion rate in the rainforests should be slower than the other habitats. We can account for this variation by fitting different speciation completion rates to different habitats, as Goldberg et al. (2011) did to associate geographic range evolution and diversification in their geographic state speciation and extinction model (GeoSSE).

For another example, Blom et al. (2016) found that, in *Cryptoblepharus,* a genus closely related to the rainbow skinks in Australia, adaptation from arboreal to rock habitat repeatedly promoted adaptive diversification, while speciation within either arboreal or rock habitat resulted in species with similar morphology. This suggests that adaptation to rock habitat drives ecological speciation, which sometimes happens so rapidly that shifts to rock habitat co-occur with speciation completion events. To account for ecological speciation, we can introduce a new parameter for how often shift to rock habitat co-occur with speciation completion event and modify ProSSE in the same way as Magnuson-Ford and Otto (2012) did to link trait changes to speciation event in their BiSSE-node enhanced state shift model (BiSSE-ness).

In summary, by using the SSE approach to model protracted speciation and extinction process, ProSSE gives accurate estimates of speciation initiation rate, extinction rate, and speciation completion rate. It is able to be extended to account for variation in speciation processes across lineages in a species group and to use not only genomic data, but other data types, such as ecological information of each tip lineage, to delimit species in a probabilistic way (Sukumaran and Knowles 2017). These properties of ProSSE make it a promising analytical tool for biologists to study speciation processes under a phylogenetic framework.

## ACKNOWLEDGEMENTS

We thank Craig Moritz’s lab for providing the unpublished tree of the Australian rainbow skinks. We thank Rampal S. Etienne for providing the performance data for PBD. We thank Craig Mortiz, Lindell Bromham, Jason Bragg, Sally Potter for valuable comments on the manuscript.

## Notes

https://github.com/huaxia1985/ProSSE

### Competing Interest Statement

The authors have declared no competing interest.

